# A mutation-resolved therapeutic atlas of NRAS-mutant melanoma reveals genotype-selective response to RAS(ON) inhibition and adaptive STAT3 survival

**DOI:** 10.64898/2026.02.18.706707

**Authors:** Sai Fung Yeung, July Xi Chen, Cherie Tsz Yiu Law, Alex Chun Ho Law, Carol Lee, Allen Ming Fai Leung, Mary Chau Po Ki, Ben Ko Chi-Bun, Wu Yue, Man Tong, Kaixi Liang, William C Cho, Michelle K. Y. Siu, Karen Kar Loen Chan, Chunning Leung, Stephen Kwok Wing Tsui

## Abstract

NRAS-mutated melanoma remains a major unmet clinical need, with no approved targeted therapy and rapid progression on standard treatment. Tri-complex RAS(ON) inhibitors such as daraxonrasib (RMC-6236) and RMC-7977 have shown early clinical activity, but the mutation-specific sensitivity landscape and adaptive resistance programs in melanoma remain undefined. To address this, we generated an isogenic 3D melanoma platform and performed a saturation mutagenesis screen across 95 NRAS missense variants (>99% of clinically recurrent variants), profiling oncogenic fitness and responses to six RAS-targeting agents in spheroids and xenografts. RMC-6236 and RMC-7977 showed the broadest activity and stratified recurrent NRAS mutants into hypersensitive (G12 variants and Q61R/K/L; ∼95% of cases), moderately sensitive (G13D/R/V; ∼4%), and resistant (G60E and Q61P; ∼1%) classes. Structural analyses supported distinct mechanisms underlying reduced susceptibility in a restricted subset of variants. In sensitive genotypes, RAS(ON) inhibition elicited an adaptive cytokine- and RTK-associated survival program converging on STAT3. Co-inhibition of STAT3 enhanced apoptosis, suppressed MYC, and induced tumor regression in NRAS-mutant melanoma models. Together, these findings define a mutation-resolved therapeutic landscape for NRAS-mutant melanoma and identify adaptive STAT3 signaling as a rational target for combination therapy.

**Statement of Translational Significance:** NRAS-mutated melanoma lacks effective targeted treatments, and clinical responses to immunotherapy are suboptimal. This study presents the first comprehensive drug sensitivity map across 95 NRAS mutations in melanoma, identifying the pan-RAS(ON) inhibitors RMC-6236 and RMC-7977 as broadly effective agents. Multiple mutants with reduced susceptibility are identified, providing mutation-informed guidance for patient selection and clinical trial stratification. Mechanistic analyses reveal that RTK/cytokine-driven STAT3 activation functions as a key survival pathway under RAS(ON) blockade, and its inhibition markedly enhances the efficacy of pan-RAS(ON) inhibitors. These findings support mutation-guided use of RAS(ON) inhibition and highlight STAT3 co-targeting as a rational strategy to strengthen and prolong therapeutic responses in NRAS-mutated melanoma.

**Highlights:**

- A functional and therapeutic atlas defines 95 recurrent and nonrecurrent NRAS missense variants in melanoma
- RMC-6236 and RMC-7977 show broad but genotype-selective activity across major NRAS mutations
- A restricted subset of recurrent NRAS mutants shows reduced susceptibility to RAS(ON) inhibition
- RAS(ON) inhibition induces an adaptive STAT3 survival program that is therapeutically targetable

## Background

NRAS mutations occur in approximately 20–30% of melanomas (1, 2, 3, 4). The current standard of care for NRAS-mutated melanoma includes immune checkpoint inhibitors, chemotherapy, and off-label use of MEK inhibitors (5, 6, 7, 8). Only 30–40% of patients exhibit initial responses, and more than half experience rapid disease progression (9, 10, 11). To date, no targeted therapies have been approved for NRAS-mutated melanoma, representing a major unmet clinical need.

Direct targeting of oncogenic RAS has recently become clinically feasible. Early evidence suggests limited repurposing of the allele-specific KRAS-G12C inhibitor sotorasib, with activity reported in rare NRAS-G12C colorectal cancers(12). This provides proof of concept but does not address the broader mutational landscape of NRAS-mutant melanoma, where the dominant hotspot is Q61. More broadly, next-generation agents such as the pan-KRAS(OFF) inhibitor BI-2865 (13) and the nucleotide-free pan-RAS inhibitor ADT-007 (14) show preclinical efficacy across multiple RAS mutants, moving beyond allele-specific targeting. However, their mutation-specific efficacy in NRAS-mutated melanoma remains unclear.

A recent advance is the emergence of tri-complex RAS(ON) inhibitors (e.g., RMC-7977, RMC-6236/daraxonrasib) (15, 16, 17). These inhibitors function as molecular glues, stabilising inactive complexes between GTP-bound RAS (mutant and wild-type) and cyclophilin A, thereby suppressing effector signaling. Clinical activity has been demonstrated in RAS-mutant solid tumors, with daraxonrasib now in a Phase III trial for pancreatic cancer (RASolute-302, NCT06625320) (16, 18). Notably, Phase Ib/II data report tumor regressions in daraxonrasib-treated NRAS-mutated melanoma, highlighting the translational potential of this class. Preclinically, combining RAS(ON) inhibition with PD-1 blockade enhances T □ cell infiltration, reprograms myeloid cells toward an antigen-presenting phenotype, and improves host survival (17, 19, 20).

Despite this progress, the mutation-specific drug efficacy landscape for NRAS-mutated melanoma remains understudied. Existing preclinical models using cell-line panels are limited by incomplete mutant representation and heterogeneous genetic backgrounds, conflating mutation-intrinsic effects with confounding variables (14, 16, 17, 21). Systematic, isogenic profiling of clinically observed NRAS variants against emerging RAS therapeutics is lacking.

Furthermore, distinct NRAS mutants are known to engage divergent signaling dependencies (22). For example, Q61 mutations show heightened dependency on RAF/MEK, whereas G12 mutations rely more on PI3K for survival. Such heterogeneity implies that both drug sensitivity and adaptive resistance to RAS(ON) inhibition are likely to be mutation-specific, yet these relationships remain poorly defined in NRAS-mutated melanoma.

To address these gaps, we used saturation mutagenesis in an isogenic NRAS-mutated melanoma model to establish 3D spheroids, enabling a high-resolution, mutation-resolved analysis of cellular fitness and drug sensitivity. We systematically profiled responses of all major NRAS mutants to six clinically relevant RAS inhibitors, defining their spectrum of activity and mapping adaptive signaling rewiring induced by RAS(ON) inhibition. These analyses indicate that most clinical NRAS mutants remain susceptible to RAS(ON) inhibition and uncover a STAT3-driven adaptive survival program that can be therapeutically targeted using combination treatment *in vitro* and *in vivo*.

## Material and methods

### Cell culture and reagents

MeWo and A375 melanoma cell lines were obtained from ATCC and maintained in MEM supplemented with 10% FBS and 1% PenStrep. sk-mel-2 melanoma cell line was obtained from ATCC and maintained in MEM supplemented with 20% FBS and 1% PenStrep. The Ba/F3 immortalized murine B cell line was a gift from Professor Zhao Hui (CUHK, SBS) and cultured in RPMI with 10% FBS, 1% PenStrep, and 10 ng/ml IL-3 (PeproTech). The Platinum-A (PLAT-A) retrovirus packaging cell line was purchased from Cell Biolabs (San Diego, CA, USA) and grown in DMEM containing 10% FBS, 1% P/S, 10 µg/mL blasticidin (Gibco, USA), and 1 µg/mL puromycin (Sigma-Aldrich, St. Louis, MO, USA). All cell lines were maintained at 37 °C in a 5% CO2 environment and confirmed to be mycoplasma-free using the VenorGeM OneStep Mycoplasma Detection Kit (Minerva Biolabs, Berlin, Germany). Authentication of the cell lines was conducted via short tandem repeat (STR) analysis (Pangenia, Toronto, ON, Canada), and results can be provided upon request. Sotorasib, Adagrasib, ADT-007, BI-2865, C188-9 were purchased from MedChemExpress (Shanghai, China), while napabucasin, RMC-6236 and RMC-7977 were obtained from ChemieTek, LLC. All drugs were prepared in DMSO.

Small interfering RNAs (siRNAs) targeting STAT3 and a non-targeting control were purchased from GenePharma (Shanghai, China); sequences are provided in Supplementary Table S2. For specificity, two independent siSTAT3 oligonucleotides were combined in transfections performed with Lipofectamine RNAiMAX (Thermo Fisher Scientific, USA) following the manufacturer’s protocol.

### Cloning, generation of stable overexpression cell lines by retroviral infection

The pMXs-puro and pMXs-EGFP-puro retroviral vectors (Cell Biolabs, USA) were used for NRAS gene delivery. The NRAS wild-type open reading frame (ORF; transcript ID: ENST00000369535.5) was subcloned into pMXs-puro by Genscript Inc. (USA) to generate pMXs-NRASWT. Site-directed mutagenesis was performed by Genscript Inc. (USA) to generate NRAS Q61K, Q61L, and Q61R mutants. Additional NRAS mutants (G12C, G13D, G13R, G13K, G60E, Q61P) were produced by site-directed mutagenesis as previously described; primer sequences are listed in Supplementary Table S2. All cloned constructs were verified by full-length sanger sequencing to confirm sequence identity.

The established constructs were then transfected into the PLAT-A cells using Lipofectamine 3000 (Invitrogen, Carlsbad, CA, USA) for 72 h. Retroviruses were filtered through a 0.45 µm polyethersulfone membrane to infect the MeWo and Ba/F3 cells for 72 h. Cells were selected with 2 µg/mL of puromycin for 7 days before downstream experiments.

### IL-3 independent growth Assay

Ba/F3-EGFP, Ba/F3-NRAS^WT^, Ba/F3-NRAS^Q61K^, Ba/F3-NRAS^Q61L^, and Ba/F3- NRAS^Q61R^ cells were maintained under identical conditions to parental Ba/F3 cells. To assess the transforming ability of NRAS mutants, cells underwent stepwise IL-3 withdrawal prior to the IL-3-independent growth assay. Cells were first adapted to 0.2 ng/mL IL-3 for 48 hours, then to 0.02 ng/mL IL-3 for an additional 48 hours. For the assay, 500 viable cells (determined by trypan blue staining) per well from each Ba/F3 stable clone were seeded in round-bottom 96-well plates. Cells were cultured in RPMI-1640 medium supplemented with 10% FBS and 1% Penicillin-Streptomycin without IL-3. Viability was quantified at indicated timepoints using the CellTiter-Glo® Luminescent Cell Viability Assay (Promega,, Madison, WI, USA) according to manufacturer protocols. Experiments were independently repeated three times.

### Spheroid formation assay

MeWo cells were seeded at 1000 cells/well in 200 µL of MEM supplemented with 10% FBS into ultra-low attachment 96-well round-bottom plates. Spheroids were cultured for 12 days in the absence of drug treatment. Images were taken using the CELENA X High Content Imaging System (Logos Biosystems, Anyang, Republic of Korea). Spheroid size was quantified by measuring cross-sectional area using ImageJ software. Spheroid cell viability was assessed using the CellTiter-Glo 3D Cell Viability Assay (Promega) according to manufacturer’s instructions at indicated time points by measuring luminescent signal. Experiments were confirmed with independent replicates.

### Drug Sensitivity Assays

#### Ba/F3 Drug Sensitivity Assays

Ba/F3–NRAS^Q61K/Q61R/Q61L^ cells were subjected to stepwise IL-3 withdrawal as previously described before drug sensitivity testing. Ba/F3-EGFP and Ba/F3-NRAS^WT^ controls were maintained in RPMI-1640 supplemented with 10% FBS and 2 ng/mL IL-3 throughout the assay. For drug screening, 1,000 viable cells (confirmed by trypan blue exclusion to avoid interference from non-viable cells) were seeded into 96-well plates in 100 μL of the appropriate growth medium. Compounds diluted in 2× concentrated medium (100 μL) were added to achieve the desired final working concentrations. Cells were incubated for 72 h at 37 °C with 5% CO □, after which viability was quantified using the CellTiter-Glo® Luminescent Cell Viability Assay (Promega). Dose–response curves and IC □ □ values were calculated relative to vehicle-treated controls (final DMSO concentration: 0.02%), which were set to 100% viability.

#### Spheroid Drug Sensitivity Assays

For 3D drug sensitivity measurement, 1,000 MeWo cells were seeded per well in 100 μL MEM supplemented with 10% FBS in ultra-low-attachment, round-bottom 96-well plates. Spheroids were allowed to form spontaneously for 72 h in a humidified incubator (37 °C, 5% CO □). Peripheral wells were filled with sterile PBS to limit evaporation. Spheroids with diameters ∼200 μm (a range that minimizes diffusion gradients and size-associated variability) were treated with compounds prepared in 2× MEM (100 μL per well). After 72 h of drug exposure, cell viability was assessed using the CellTiter-Glo® 3D assay (Promega) following the manufacturer’s protocol. All experiments were confirmed with three independent replicates.

#### Flow Cytometry Analysis

After completion of each indicated treatment, cells were stained with Annexin V–FITC and propidium iodide (BD Biosciences, San Jose, CA, USA) according to the manufacturer’s instructions and analyzed on a BD FACSymphony A5.2 SORP flow cytometer (BD Biosciences, San Jose, CA, USA).

#### Animal Experiments

All animal experiments were carried out in accordance with protocols approved by the Animal Experimentation Ethics Committee of the Chinese University of Hong Kong (CUHK; reference and consent number: 22-257-MIS). Female nude mice (4–6 weeks old) were bred and housed in the Laboratory Animal Services Centre, CUHK. Detailed *in vivo* procedures were summarized in supplementary materials.

#### Statistical Analysis

Statistical analyses were performed using GraphPad Prism (version 10.5.0; serial number GPS-2505185-TIT3-F100A; RRID: SCR_002798). Data are presented as mean ± SEM unless otherwise specified. A p-value < 0.05 was considered statistically significant. Group comparisons were performed using one-way ANOVA followed by Šidák’s post hoc multiple-comparisons test. For clarity, p-values reported in the main text reflect comparisons between the wild-type and the corresponding mutant groups only.

Detailed experimental methods, including cell line generation, retroviral transduction, saturation mutagenesis, drug screening, and structural modeling, are provided in the Supplementary Materials.

## Results

### Prevalence, oncogenicity and signaling of common NRAS mutations in a melanoma model

To assess the potential use of common RAS-targeted agents in NRAS-mutant melanoma, we first characterized the NRAS mutation landscape. Analysis of combined melanoma datasets (N = 878) showed that NRAS mutations occur predominantly at the P-loop (G12/G13: 8.8%) and the Switch II domain (A59/Q61: ∼83.8%), consistent with COSMIC frequencies (Fig. 1A). The most prevalent variants were Q61R (32.5%), Q61K (29.7%), and Q61L (16.3%) with additional less frequent mutations at these codons. To compare their functional output, we generated isogenic Ba/F3 and MeWo models expressing GFP, NRAS^WT^, NRAS^Q61R^, NRAS^Q61K^, or NRAS^Q61L^. Ba/F3 cells (IL-3–dependent murine pro-B cells) allow assessment of IL-3–independent growth driven by oncogenic NRAS signaling, wheras MeWo cells allow the evaluation of enhanced 3D proliferation and tumorigenicity in a melanoma background. Because the MeWo cell line is NF1-null, it exhibits constitutive baseline RAS/RAF activation. Because MeWo cells are NF1-null, they lack a major GAP-mediated negative regulatory input and therefore exist in a primed RAS/RAF signaling state. This context permits exogenous NRAS mutants to further amplify downstream signaling above the NF1-null baseline.

**Figure 1.**
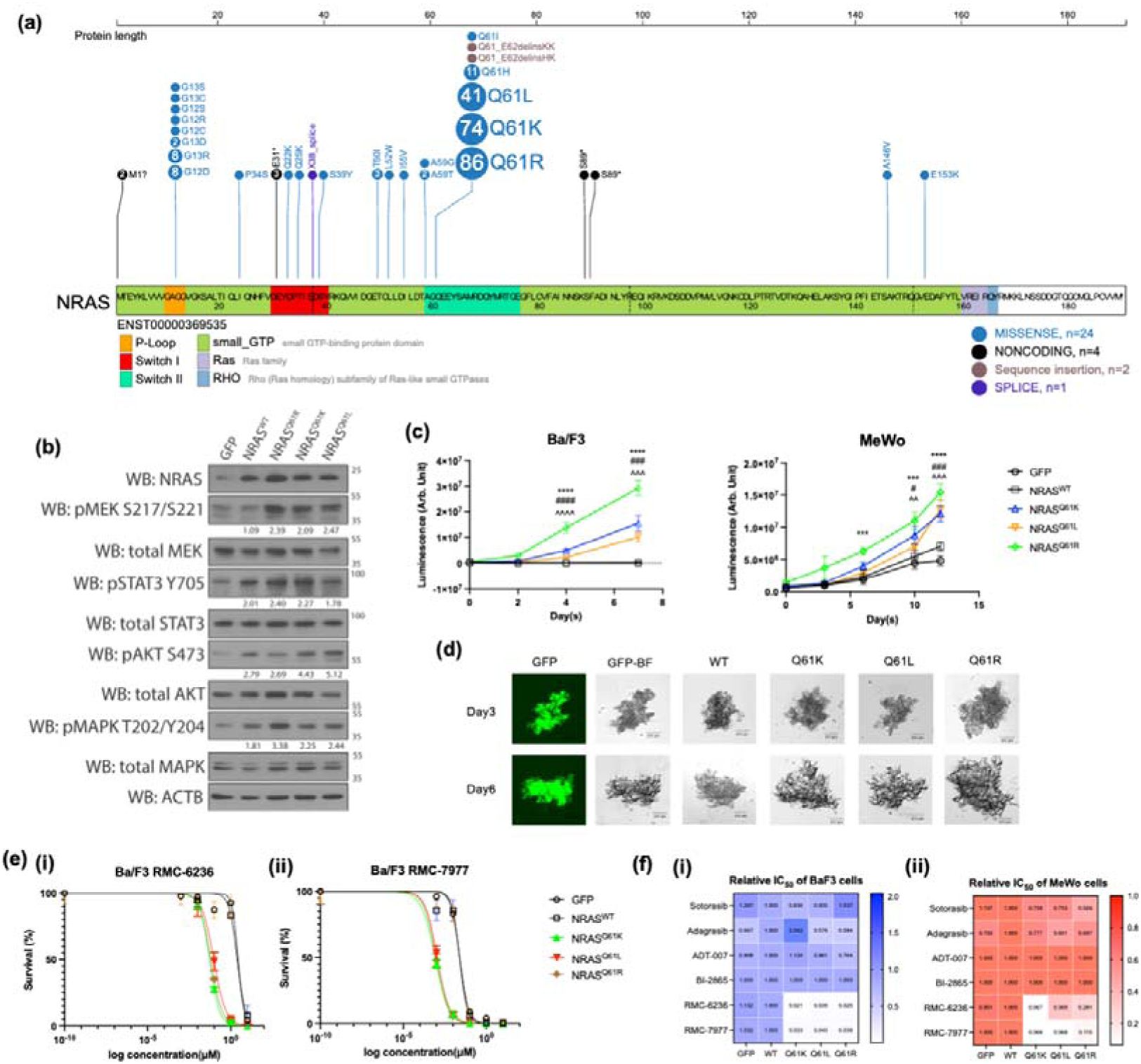
NRAS mutation hotspots in melanoma and their differential sensitivity to active RAS(ON) inhibitors RMC-6236 and RMC-7977. (A) Distribution of NRAS mutations in combined dataset from TGEN, Genome Res 2017(1), MSK, Clin Cancer Res 2021(2), and DFCI, Nature Medicine 2019(3), data visualized using ProteinPaint plot from the GDC Genome Browser. (B) Western blot analysis of MAPK, AKT, and STAT3 signaling in isogenic MeWo cells expressing GFP (control), NRAS^WT^, NRAS^Q61R^, NRAS^Q61K^, or NRAS^Q61L^. Signal intensities were quantified by densitometry using ImageJ normalized using GFP. Data confirmed with independent experiments. (C) CellTiter-Glo proliferation assays in (left) Ba/F3 cells cultured without IL-3 to assess IL-3–independent oncogenic fitness over 7 days, and (right) MeWo 3D spheroids growth monitored over 12 days (n = 6). Statistical comparisons: * = NRAS^Q61R^ vs. NRAS^WT^; # = NRAS^Q61K^ vs. NRAS^WT^; ^ = NRAS^Q61L^ vs. NRAS^WT^. *p < 0.05, **p < 0.01, ***p < 0.001, ****p < 0.0001; two-way ANOVA with Sidak’s multiple-comparisons test. For clarity, only wild-type vs. mutant comparisons are reported. Data confirmed with independent experiments. (D) Representative GFP fluorescence and bright-field image of isogenic MeWo cells expressing each NRAS variant, demonstrating relative proliferative capacity. (E) Dose–response curves for RMC-6236 and RMC-7977 (72 h) in Ba/F3 cells (1,000 cells/well) expressing GFP, NRAS^WT^, or NRAS^Q61R/K/L^. GFP and NRAS^WT^ cells were maintained in 2 ng/mL IL-3, whereas NRAS-mutant cells were cultured in IL-3–free RPMI-1640 with 10% FBS to assess IL-3-independent survival and inhibitor sensitivity. CellTiter-Glo 3D was used for viability assay. Data confirmed with independent experiments. (F) Heatmap of relative half-maximal inhibitory concentrations (IC □ □) for six RAS-targeting agents (sotorasib, adagrasib, ADT-007, BI-2865, RMC-6236, RMC-7977) measured at 72 h in Ba/F3 cells (IL-3–free for mutants; IL-3 supplemented for GFP/WT) and in pre-formed MeWo spheroids treated for 72 h. Each IC □ □value for a mutant–drug pair is expressed relative to the corresponding NRAS^WT^ IC □ □for that drug. Data confirmed with three independent experiments.

In MeWo cells, NRAS^Q61R/K/L^ significantly increased 3D spheroid growth over 12 days compared with NRAS^WT^ and GFP controls (p < 0.001; Fig. 1C, D; Fig. S1). In Ba/F3 assays, the same variants conferred robust IL-3–independent proliferation over 7 days (p < 0.001; Fig. 1C). NRAS^Q61R^ exhibited the strongest oncogenic activity (∼3-fold induction), whereas NRAS^Q61K^ and NRAS^Q61L^ showed approximately half the activity of NRAS^Q61R^. In MeWo cells, all variants elevated p-STAT3, p-MEK, and p-MAPK relative to NRAS^WT^, with NRAS^Q61K/L^ also inducing stronger p-AKT (Fig. 1B).

We next evaluated drug sensitivity to RAS-targeting agents to validate our platform. NRAS^Q61R/K/L^ showed marked hypersensitivity to the RAS(ON) inhibitors RMC-6236 and RMC-7977 (Fig. 1E–F), whereas KRAS-G12C inhibitors (sotorasib, adagrasib, BI-2865) and ADT-007 showed no consistent activity pattern across these mutants (Fig. 1F). In MeWo cells, NRAS^Q61K/L^ displayed modestly reduced sensitivity to RMC-6236 compared with NRAS^Q61R^. Together, these results demonstrate substantial biochemical heterogeneity among common NRAS Q61 variants and highlight mutant-specific drug response.

### Comprehensive Functional Profiling and Drug Sensitivity Mapping of NRAS Missense Mutations in Melanoma

To capture the functional diversity of NRAS mutations beyond these common variants, we generated a saturation mutagenesis library of 95 missense NRAS variants derived from an NNK codon library spanning 160 possible codon-level variants across five hotspot residues at G12, G13, A59, G60 and Q61. This panel covers the most frequently reported mutations in the P-loop and switch-II domain and enabled systematic profiling of oncogenic fitness and drug sensitivity. Fitness profiling using pooled competitive 3D spheroid assays for 9 days and xenograft competitive growth assays for 16 days (Fig. 2A, B; Fig. S4) showed that G13- and Q61-containing mutants dominated the top enriched variants, including the recurrent NRAS^G13R^ and NRAS^Q61H/L^. NRAS^Q61R^ consistently displayed a strong fitness advantage *in vitro* (2.1-fold) and *in vivo* (1.46-fold), whereas NRAS^Q61K^ exhibited high *in vitro* fitness (2.24-fold) but no *in vivo* advantage (0.79-fold).

**Figure 2.**
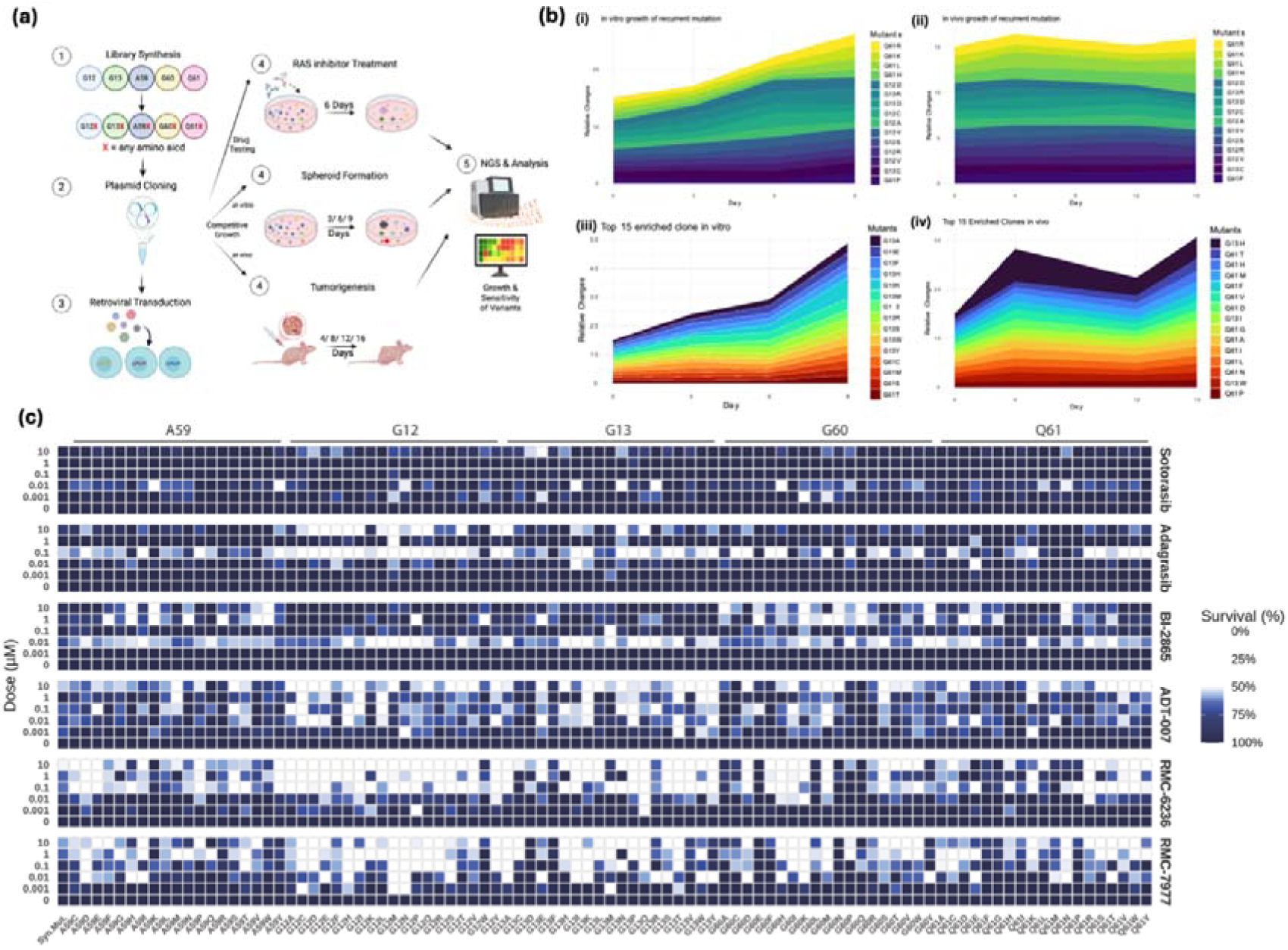
Saturation mutagenesis mapping of NRAS variant fitness and RAS(ON) inhibitor sensitivity in isogenic MeWo melanoma cells. **(A)** Schematic overview of the NRAS saturation mutagenesis workflow, competitive growth profiling, and inhibitor-sensitivity screening. **(B)** Temporal dynamics of NRAS variant abundance relative to synonymous controls during pooled competitive assays. MeWo cells expressing all 95 missense mutants were cultured either as 3D spheroids in low-attachment plates with MEM + 10% FBS (i and iii) or injected subcutaneously into nu/J mice (ii and iv). For each time point, variant frequencies were averaged across biological replicates (n = 4) and normalized to day-0 values. Colors denote individual variant trajectories. **(C)** Drug-response profiling of pooled NRAS mutants under treatment with sotorasib, adagrasib, BI-2865, ADT-007, RMC-6236, and RMC-7977. Read counts for each mutant abundance in inhibitor-treated pools (day 6) were normalized to DMSO controls. Relative viability (%) is color-coded according to the scale shown.

We next profiled the drug sensitivities of all variants against the six RAS-targeting agents described above. Resistance or sensitization was defined by differential mutant abundance relative to NRAS^syn.mut^ in inhibitor- versus DMSO-treated conditions at Day 6. Consistent with Fig. 1F and previous reports, NRAS^Q61R/K/L^ mutants showed strong sensitivity to the tri-complex inhibitors RMC-6236 and RMC-7977 but responded only weakly to other agents (Fig. 2C). Approximately 80% of G12/G13 mutants were also highly sensitive to these two inhibitors. Among the 23 clinically recurrent NRAS mutations, four (G13D, Q61P, G13R, G60E) were resistant to both tri-complex inhibitors, NRAS^G13V^ showed preferential resistance to RMC-7977, and the remaining 18 (78.3%, mutation-count based) were sensitive (Fig. 2C; Fig. S6). These mutations, when weighted by clinical mutation frequency, these sensitive classes accounted for ∼95% of recurrent NRAS-mutant melanoma cases. Notably, many sensitizing G13 variants (G13K/E/W/S/Y) overlapped with the top fitness-enhancing mutants identified in the competitive growth assays.

In contrast to the clear profiles observed for RMC-6236/RMC-7977, no coherent sensitivity profile emerged for the KRAS-G12C inhibitors sotorasib or adagrasib. Although a prior study reported sotorasib activity against NRAS^G12C^(12), we observed only minimal effects in our melanoma model (Fig. 2C), suggesting a melanoma-specific resistance mechanism, while adagrasib showed no consistent mutant-dependent response similarly. By comparison, the pan-RAS inhibitor ADT-007 exhibited selective activity toward a subset of G12/G13/G60 mutants, including the clinically recurrent G12C mutant, with 15 variants showing increased sensitivity relative to NRAS^syn.mut^. BI-2865 also demonstrated inhibitory activity against a subset of G60 mutants. Structural modeling indicated potential non-covalent interactions underlying this selective activity (Table S4; Fig. S8–S9), and several ADT-007 responsive G13 mutants overlapped with top-enriched variants from the fitness screens, particularly NRAS^G13K/E/W/S/Y^ (Fig. 2C; Fig. S6). However, this narrow sensitivity profile may reflect intrinsic targeting properties of ADT-007 (Fig. S9; Table S4), or possible UGT-mediated drug inactivation in MeWo cells (18).

In short, these data show that RAS(ON) inhibitors RMC-6236 and RMC-7977 potently target NRAS-mutant melanoma, with ∼95% of clinically recurrent cases classified as sensitive in our atlas (Table S5). ADT-007 and BI-2865 provide additional coverage for rarer mutants, whereas RAS(ON) inhibitor resistance is concentrated in a restricted subset of clinically observed variants, including NRAS^Q61P^, NRAS^G13D/V/R^, and NRAS^G60E^ (Table S5).

### Structural Basis of Altered Drug Sensitivity in NRAS Mutants

To validate our findings with RMC-6236/RMC-7977 from the saturated mutagenesis screen, we generated individual MeWo clones expressing representative NRAS mutations (G12C, G13D/R/V, G60E, Q61P). In 3D spheroid assays, most mutants induced ∼2-fold growth compared with NRAS^WT^ over 8 days, except NRAS^G60E^, which showed weaker activity (1.1-fold) (Fig. 3B, 3C). Drug-response assays faithfully recapitulated pooled-screen patterns: NRAS^G12C^ was strongly sensitized to tri-complex inhibition (14.53-fold and 5.6-fold lower IC □ □ for RMC-6236 and RMC-7977), mirroring canonical Q61 sensitizers (Fig. 1F; Fig. 3D). NRAS^G13D/R/V^ variants exhibited only modest IC □ □ shifts (0.26–2-fold), whereas NRAS^G60E^ and NRAS^Q61P^ remained resistant (1.3–2.7-fold increased IC □ □) (Fig. 3D; Fig. S11).

**Figure 3.**
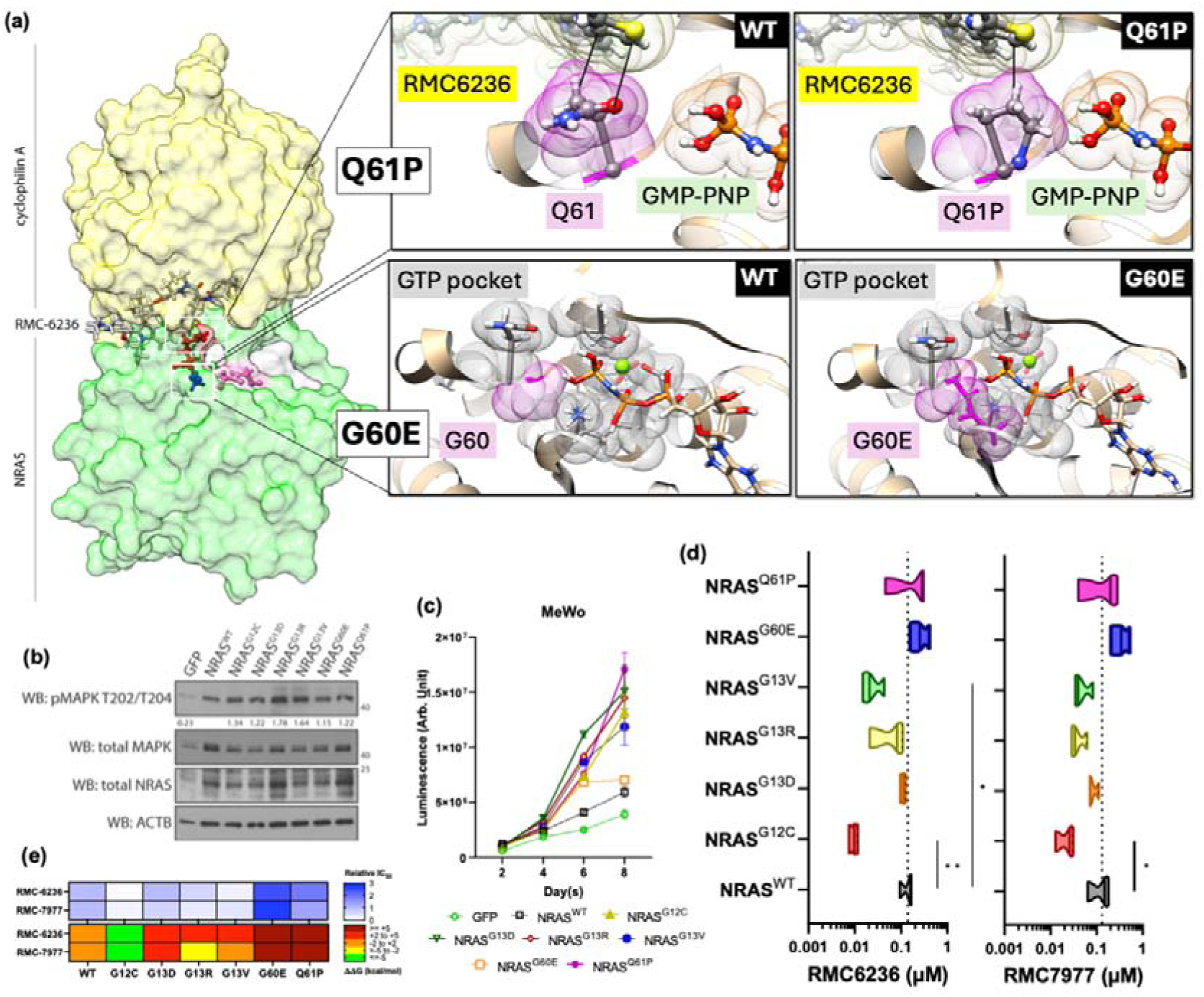
Cellular and structural characterization of NRAS mutants with reduced susceptibility to RAS(ON) inhibitors. **(A)** Structural analyses of intrinsic resistance mutants. (Left) Overall crystal structure of the Cyclophilin A–NRAS–RMC-6236 tri-complex, highlighting key residues: Q61 (red), G60 (blue), and G12/G13 (pink). (Right) Modeled co-crystal structures showing van der Waals surfaces of residues 60/61, RMC-6236, and GMP-PNP. In NRAS^WT^, Q61 maintains optimal molecular distances with both RMC-6236 and GMP-PNP without steric interference. The Q61P substitution introduces steric clashes between Pro61 and the thiazole moiety of RMC-6236 (HG2–C9) and with GMP-PNP (HD3–HO2G, C10–HG2, CD–H2OG), disrupting inhibitor–RAS contacts. The G60E substitution introduces steric clashes between Glu60 and the GTP-binding pocket. **(B)** Western blot confirming expression of selected NRAS variants and associated signaling changes. **(C)** Oncogenic fitness of selected NRAS mutants in MeWo isogenic clones. 3D spheroid growth was monitored over 8 days. Findings were reproducible across independent experiments. **(D)** IC_50_ distributions of selected NRAS mutants from the saturation mutagenesis panel treated with RMC-6236 or RMC-7977. Statistical significance relative to NRAS^WT^ was determined by unpaired t-test from three independent experiments (*p < 0.05, **p < 0.01). **(E)** Heatmap showing the correlation between relative drug sensitivity (IC_50_) and predicted thermostability changes (ΔΔ G, kcal/mol) for each mutation, modeled using the structure (PDB: 9BG0).

To understand the mechanisms underlying the mutant-specific drug responses, we modeled NRAS–PPIA–inhibitor tricomplexes using available co-crystal structures (PDB: 9BG0, 8TBI) (15, 17). NRAS^Q61P^ destabilized the tricomplex by abolishing the Q61-mediated polar contact with the inhibitor’s S1 atom and disrupting hydrophobic interactions with the RMC-6236 thiazole moiety. NRAS^Q61P^ also introduced steric clashes with GMP-PNP and reduced hydrophobic packing against the C10 group of RMC-6236 (Fig. 3A). Distinct structural incompatibilities induced by NRAS^G60E^ and NRAS^Q61P^ with RMC-7977 are shown in Fig. S7.

In contrast, NRAS^G13D/R/V^ are located distal to the tricomplex interface and produce no direct atomic clashes (Fig. 3A). However, *in silico* analyses predicted that these substitutions generally reduce binding affinity and weaken tricomplex stability, with the notable exception of NRAS^G13R^, which was uniquely predicted to decrease thermal stability in the presence of either RMC-6236 or RMC-7977 (Fig. 3E). For RMC-6236, the predicted Δstability values (kcal/mol) were +3.569 (G13D), +2.483 (G13R), and +2.612 (G13V). For RMC-7977, the values were +2.794 (G13D), −1.991 (G13R), and +1.424 (G13V).

Collectively, these analyses reveal distinct sensitivity profiles among clinically recurrent NRAS variants. Relative to wild-type NRAS, our data define three functional classes: hypersensitive (G12C, Q61R/K/L; ∼95% of cases; >5-fold IC □ □ reduction), moderately sensitive (G13D/R/V; ∼4% of cases; modest IC □ □ shifts), and resistant (G60E, Q61P; ∼1% of cases; marked IC □ □ increase).

### STAT3 activation is a mutation-enhanced adaptive survival mechanism under RAS(ON) inhibition

Melanoma frequently develops resistance to monotherapy, leading to relapse and limited treatment options (23, 24). To improve the depth and durability of responses, we examined adaptive signaling pathways engaged upon RAS inhibition in hypersensitive mutants to identify potential combination strategies. Specifically, we profiled four major survival nodes (MAPK, PI3K/AKT, TOR/p70S6K, and JAK–STAT) following short-term treatment with different inhibitors in a dose-dependent manner.

Western blot analysis indicated that neither the KRAS inhibitors (sotorasib, adagrasib, BI-2865) nor ADT-007 induced coherent signaling changes across NRAS^WT^ and NRAS^Q61^ mutant lines (Fig. 4A; Fig. S2). Only RMC-6236 and RMC-7977 uniformly suppressed p-MAPK across all backgrounds. MAPK inhibition plateaued in NRAS^WT/Q61K/L^ but was complete in NRAS^Q61R^. Both inhibitors induced strong activation of p-STAT3 Y705, with the highest induction in NRAS^Q61R/L^ (Fig. 4A; Fig. S2). Prolonged exposure (12–24 h) further amplified STAT3 activity in the MeWo overexpression system, where NRAS^Q61R/K/L^ variants exhibited substantially higher induction than NRAS^WT^ cells (Fig. S13B), suggesting a mutation-enhanced adaptive response. Importantly, this trend was mirrored in endogenous models, the NRAS^Q61R^ sk-mel-2 line showed a 1.3–1.5-fold increase in p-STAT3 levels over a 0–8 hour treatment time course, whereas no such upregulation was observed in endogenous NRAS^WT^ lines, including parental MeWo and A375 cells (Fig. 4B).

**Figure 4.**
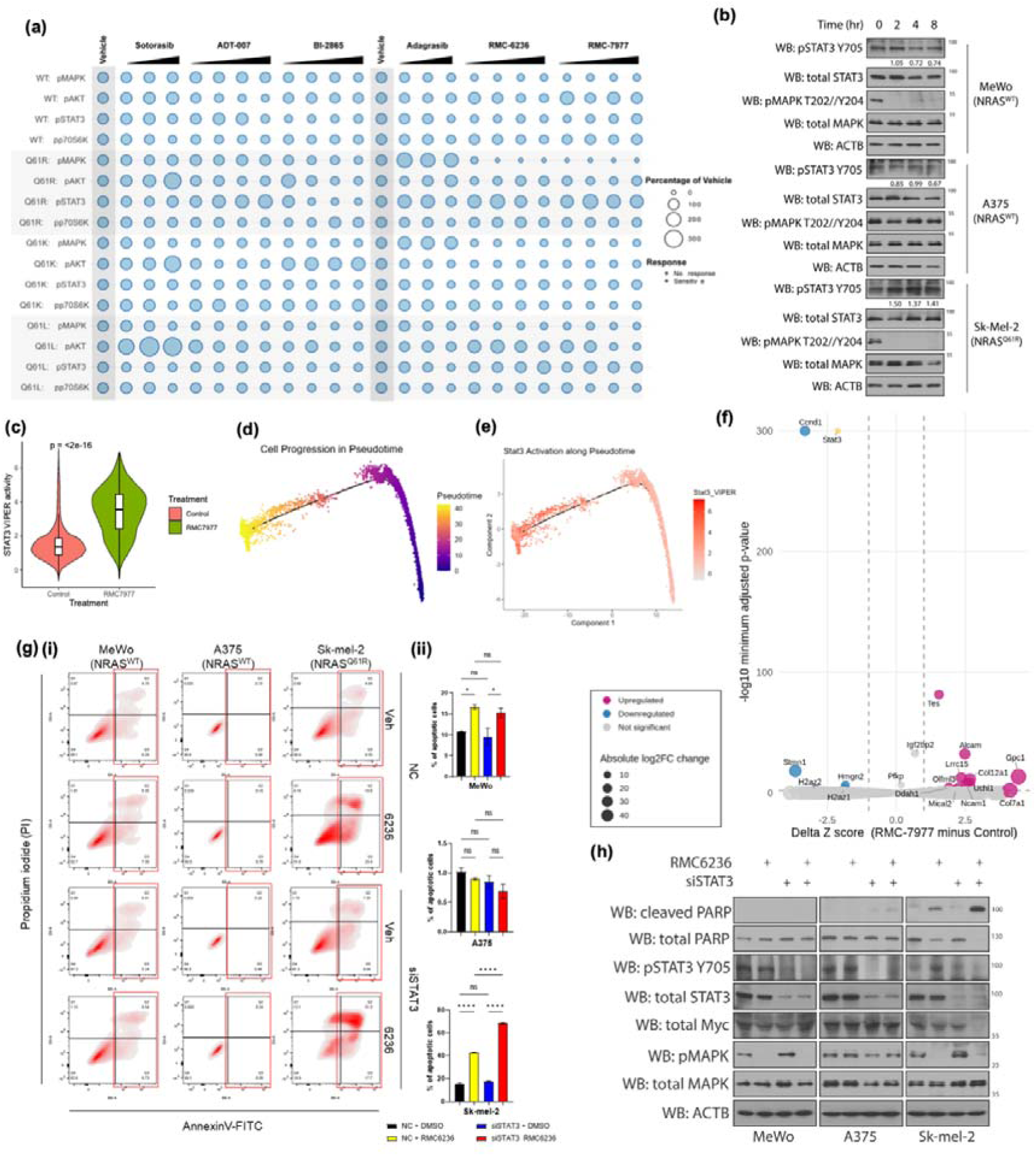
STAT3 is required for survival of melanoma cells treated with RAS(ON) inhibitors. (a) Signaling responses to RAS(ON) inhibition in isogenic MeWo cells. The bubble plot depicts differential signaling responses across *NRAS* variants following 8-hour treatment with the indicated inhibitors. Signaling was assessed by Western blot, with band intensities quantified by densitometry and normalized to vehicle controls (set to 100%). Bubble size represents normalized signaling levels. Cells were treated with dose ranges specific to each inhibitor class: sotorasib and adagrasib (0.1, 1, 10 µM); ADT-007, BI-2865, RMC-6236, and RMC-7977 (0.01, 0.1, 1, 10 µM). (b) Western blot analysis of phospho-STAT3 (Tyr705) in NRAS^WT^ (MeWo, A375) and NRAS^Q61R^ (sk-mel-2) cells treated with 1 µM (WT) or 0.1 µM (Q61R) RMC-6236 for 0, 2, 4, and 8 hours. (c) Violin plot comparing STAT3 VIPER scores (inferred transcription factor activity) between vehicle-control and RMC-7977 treated malignant melanoma cells from the GSE300712 cohort. (d) Trajectory plot demonstrating the Monocle-inferred pseudotime evolution of malignant cells isolated from vehicle-control and RMC-7977 treated O.SUMMER13 mouse melanoma tumors obtained from GSE300712. (e) Overlay of STAT3 VIPER activity scores mapped across the pseudotime trajectory defined in (d). (f) Volcano plot illustrating the *scTenifoldKnk* virtual knockout effect of *Stat3* in control versus RMC-7977 treated malignant cells from GSE300712. Z-scores represent the statistical significance of the manifold displacement for each gene upon virtual *Stat3* deletion. The differential Z-score identifies an induced dependency (red) in genes involved in RTK-scaffolding and adhesion proteins versus a loss of dependency (blue) in cell-cycle regulators. (g) Analysis of apoptosis induced by RAS(ON) inhibitors in combination with STAT3 knockdown. (i) Representative flow cytometry density plots of Annexin V-FITC/PI staining in NRAS^WT^ (MeWo and A375) and NRAS^Q61R^ (sk-mel-2) cells after 16-hour treatment with RMC-6236; red gates indicate the apoptotic population. (ii) Quantitative analysis of total apoptotic cells (mean ± SD, n=3). Statistical significance was determined by two-way ANOVA followed by a post-hoc multiple comparisons test (∗*p*<0.05,∗∗*p*<0.01,∗∗∗*p*<0.001,∗∗∗∗*p*<0.0001, ns = no significant). (h) Western blot analysis of downstream signaling and the apoptotic marker cleaved PARP in melanoma cells transfected with non-targeting (siNC) or *STAT3*-targeting (siSTAT3) siRNA and treated with RMC-6236 (1 µM for MeWo/A375; 0.1 µM for sk-mel-2).

To provide an orthogonal validation of these signaling dynamics within the broader transcriptomic landscape, we interrogated single-cell RNA sequencing (scRNA-seq) data from the GSE300712 mice melanoma cohort. This dataset encompasses vehicle-control and RMC-7977-treated tumors derived from the O.SUMMER13 model. VIPER-based inference of transcription factor activity further confirmed a significant elevation of STAT3 activity in malignant cells surviving RMC7977 treatment compared to control (Fig. 4C). Pseudotime reconstruction of the malignant cell population revealed a continuous evolutionary trajectory. Vehicle-control cells occupied the early, treatment-naïve states and progressed toward a distinct, RMC-7977-enriched adaptive terminus (Fig. 4D). Notably, when mapped across the pseudotime continuum, STAT3 activity exhibited a progressive increase as cells transitioned from the early control states toward the late drug-adapted state. This temporal correlation identifies STAT3 activation as a core feature of the transcriptional adaptation required for drug-persistence under RMC-7977 inhibition (Fig. 4E).

To further examine the functional role of STAT3 within this treatment-adapted regulatory state, we performed virtual knockout analysis. Using *scTenifoldKnk*, we simulated the loss of Stat3 within the inferred gene regulatory networks. In control cells, Stat3 was primarily linked to canonical proliferative modules, including regulators of cell-cycle progression (e.g. CCND1) as expected. In contrast, Stat3 deletion in RMC-7977-treated cells preferentially disrupted genes associated with cellular architecture, adhesion, and receptor-associated signaling networks (e.g. Tes, Gpc1) (Fig. 4F). These results indicate that under RAS(ON) inhibition, Stat3 becomes increasingly coupled to transcriptional programs that support cellular adaptation and survival to achieve a drug-persistent state.

We next tested whether STAT3 activity functionally contributes to melanoma cell survival under RAS inhibition. To determine whether STAT3 supports survival during RAS blockade, we depleted STAT3 via siRNA-mediated knockdown followed by RMC-6236 treatment in NRAS^WT^ (MeWo, A375^BRAFV600E^) and NRAS^Q61R^ (sk-mel-2) cells (Fig. 4G). STAT3 knockdown was confirmed by Western blot, which demonstrated effective suppression of p-STAT3 (Tyr705) levels (Fig. 4H). In sk-mel-2 cells, the apoptotic population (Annexin V^+^) increased significantly from 42.63% to 68.47% (1.61-fold increase; p<0.0001) (Fig. 4G). Similarly, within the MeWo overexpression model, STAT3 depletion with RMC-6236/-7977 in NRAS^WT^ cells resulted in a baseline apoptotic population of 51.56%, which further increased to approximately 70–80% in NRAS^Q61R/K/L^ variants, representing a 1.4–1.5-fold enhancement (p<0.0001) (Fig. S10 and S13).

Consistent with the apoptotic assays, Western blot analysis revealed that combined STAT3 depletion and RAS(ON) inhibition (RMC-6236 or RMC-7977) markedly increased cleaved PARP levels specifically in NRAS-mutant models, including sk-mel-2 (endogenous NRAS^Q61R^) and MeWo cells overexpressing NRAS^Q61R/K/L^ variants (Fig. 4D and S13). In contrast, parental MeWo (NRAS^WT^), A375, and MeWo cells overexpressing NRAS^WT^ displayed only marginal increases in cleaved PARP under the same co-inhibitory conditions. A similar divergence was observed in the regulation of the key survival effector, MYC. While RAS(ON) inhibitors alone achieved only partial MYC suppression, their combination with siSTAT3 triggered near-complete MYC ablation in both the endogenous sk-mel-2 and NRAS^Q61R/K/L^-overexpressing MeWo systems. Conversely, MYC remained clearly detectable in NRAS^WT^ models, including parental MeWo and A375 cells (Fig. 4H and S13). These findings indicate that complete MYC loss represents a mutant-specific vulnerability revealed only upon dual RAS and STAT3 blockade. Furthermore, these data suggest that while NRAS^WT^ cells are less dependent on the STAT3/MYC axis under RAS inhibition, NRAS-mutant cells develop an acute reliance on this compensatory circuit for survival.

### Compensatory cytokine and RTK feedback programs accompany adaptive STAT3 activation following RAS(ON) inhibition

Having established STAT3 as an adaptive survival node under RAS(ON) inhibitor blockade, we next asked which upstream programs may reactivate this pathway. To address this, we re-examined the scRNA-seq data from Control and RMC-7977-treated O.SUMMER13 murine melanoma tumors. UMAP projection showed that malignant cells from RMC-7977-treated tumors occupied a transcriptionally distinct cluster relative to control tumors, indicating that RAS(ON) inhibition induces a discrete adaptive tumor-cell state (Fig. 5A). Projection of PROGENy scores onto the malignant-cell compartment further demonstrated increased inferred STAT3 activity in the RMC-7977-treated cluster compared with control cells (Fig. 5C). The STAT3-associated transcriptional dynamics along a Monocle-inferred pseudotime trajectory based on the top 50 STAT3 target genes resolved into three major cluster. First, a early declining cluster enriched for baseline proliferative-state genes, such as Myc, Ccnd1, and Cdk4. Second, an induced immediate-early inflammatory/cytokine-responsive cluster containing Il6, Il6st, and Socs3, etc, consistent with a cytokine-responsive inflammatory program. And a late cluster containing Hif1a, Ccl2, Twist etc, consistent with a broader adaptive survival and remodeling state that also incorporates inflammatory mediators (Fig. 5B). Together, these data suggest that RAS(ON) inhibition shifts melanoma cells away from a baseline proliferative state toward a cytokine-responsive, stress-adaptive phenotype.

**Figure 5.**
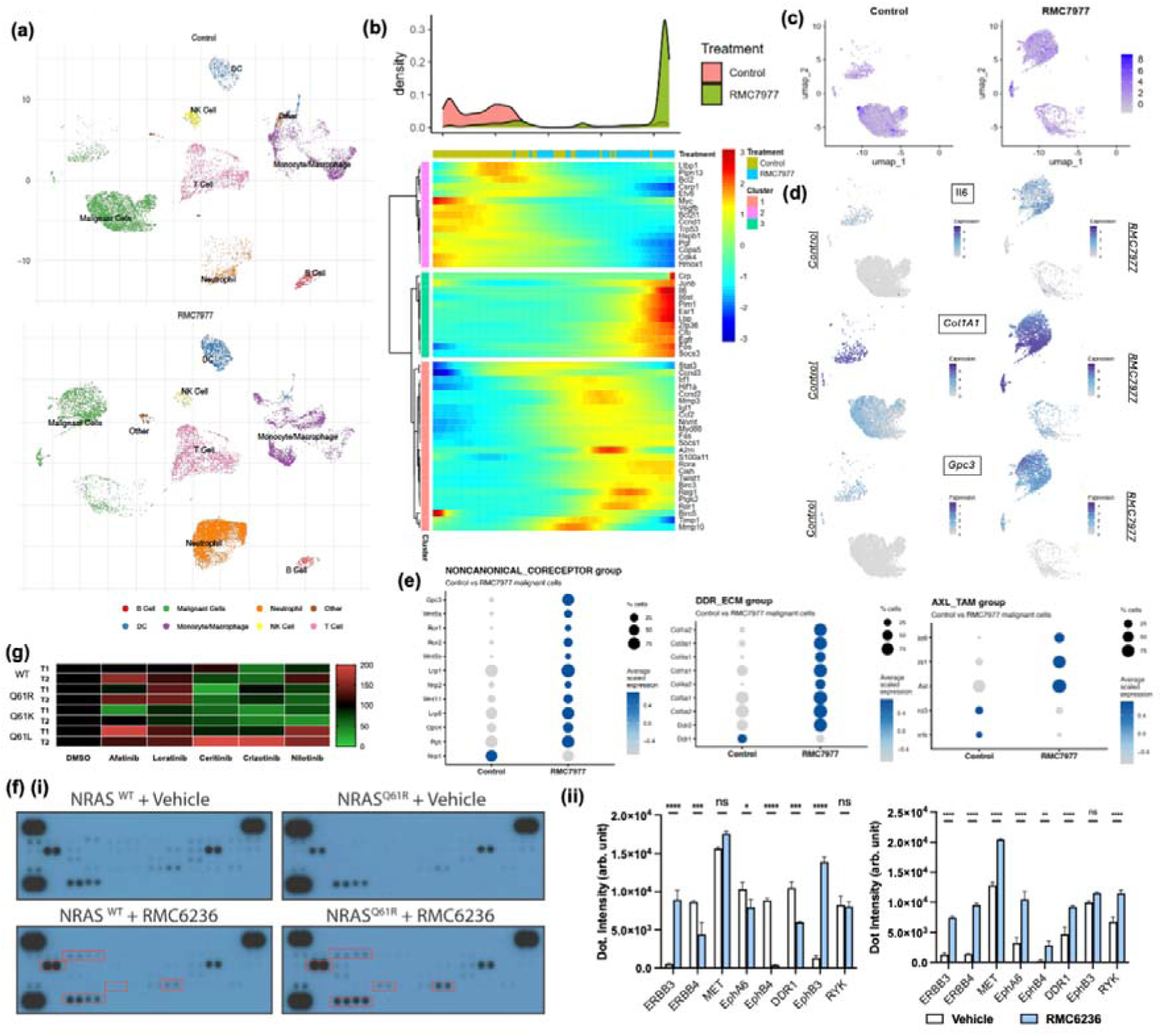
STAT3-driven adaptive survival programs are activated via compensatory RTK feedback following RAS(ON) inhibition. (a) UMAP projection illustrating the cell type distribution within the GSE300712 atlas, comparing vehicle control and RMC-7977-treated murine melanoma tumors. (b) Density plot (upper) and heatmap (lower) displaying the expression kinetics of the top 50 STAT3 target genes along a Monocle-inferred pseudotime trajectory using scRNA-seq data from the GSE300712 malignant cell population. (c) Feature plots demonstrating the spatial distribution of PROGENy scores for estimated STAT3 activity in malignant cells under vehicle control or RMC-7977 treatment conditions. (d) Feature plots identifying significantly upregulated expression of specific ligands, cytokines, and the non-canonical RTK co-receptor *Gpc3* in RMC-7977-treated populations compared to Control. (e) (e) Dot plot summarizing the expression of diverse RTK signaling components, including AXL, ECM-related receptors, and surface ligand co-receptors extracted from the GSE300712 malignant cell clusters. (f) Analysis of compensatory RTK signaling following RAS(ON) blockade: (i) Representative phospho-receptor tyrosine kinase (RTK) array blots of MeWo NRAS^WT/Q61R^ cells treated with 1 µM RMC-6236 or vehicle for 16 hours. (ii) Densitometric quantification of significantly altered phospho-RTK signals. Data represent mean ± SD. Statistical significance was determined by two-way ANOVA (*p<0.05, **p<0.01, ***p<0.001, ****p<0.0001). (g) Heatmap depicting relative STAT3 phosphorylation levels in isogenic MeWo cells co-treated with 1 µM RMC-6236 and 1 µM of indicated RTK inhibitors (lorlatinib, afatinib, ceritinib, nilotinib, and crizotinib) for 24 hours. Band intensities were quantified via densitometry and normalized to DMSO-treated controls (100%).

On this basis, we next examined whether treated cells had also acquired a cytokine-and RTK-permissive state (25). Feature-plot analysis revealed increased expression of ligands, cytokines, and non-canonical co-receptors in the treated population, including Il6, Hgf, and Gpc3 (Fig. 5D). In parallel, dot-plot analysis showed broader upregulation of RTK-associated signaling components across treated malignant clusters, encompassing HER-family receptors, the MET axis, AXL-related signaling, extracellular matrix-linked receptor programs, and additional classical RTK groups (Fig. 5E, S15). These transcriptional changes suggest that RAS(ON) blockade induces a growth factor- and cytokine-rich adaptive state that is poised to reinforce STAT3 signaling through convergent inflammatory and RTK-linked inputs.

To obtain biochemical evidence for compensatory receptor activation, we next performed phospho-RTK array analysis in MeWo NRAS^WT/Q61R^ cells treated with RMC-6236. Multiple RTKs displayed increased phosphorylation following RAS(ON) blockade, with the strongest induction observed for ERBB3, ERBB4, and EphA6, together with additional activation of MET, DDR1, RYK, and other Eph-family members (Fig. 5F). These findings provide supportive information for the adaptive receptor program suggested by the single-cell data and indicate that RAS(ON) inhibition elicits a broad compensatory signaling response rather than dependence on a single upstream receptor.

To test whether this receptor rewiring contributes functionally to STAT3 reactivation, we co-treated MeWo cells with RMC-6236 and a panel of RTK inhibitors. Ceritinib and nilotinib more effectively suppressed STAT3 phosphorylation than afatinib, crizotinib, or lorlatinib (Fig. 5G), arguing that STAT3 reactivation is most likely driven by a redundant multi-RTK network rather than one dominant receptor. Collectively, these data support a model in which RAS(ON) inhibition reshapes melanoma cells into a cytokine- and growth factor-rich adaptive state in which inflammatory signaling and compensatory RTK activation converge on STAT3 to preserve residual tumor-cell survival under drug pressure.

### STAT3 inhibitor enhances the therapeutic efficacy of RAS(ON) inhibitors in NRAS-mutated melanoma

To translate our mechanistic findings into a therapeutic strategy, we investigated whether the induced STAT3 dependency constitutes a targetable vulnerability in NRAS-mutant melanoma. We performed viability-based synergy assays using a 1:1 dilution series of RMC-6236 and the STAT3 inhibitor C188-9. Results revealed synergy in endogenous NRAS-mutant sk-mel-2 cells (HSA: 12.23 ± 0.428), whereas NRAS^WT^ parental lines MeWo (-1.827 ± 1.256) and A375 (-1.180 ± 0.777) exhibited slight antagonism (Fig. 6C). To rule out the potential off-target effect, a second STAT3 inhibitor Napabucasin is included. Consistently, napabucasin recapitulated these findings in the MeWo overexpression (OE) system. NRAS^Q61R^ (10.87 ± 1.80) and NRAS^Q61K^ (11.26 ± 0.92) showed robust synergy, NRAS^Q61L^ showed moderate synergy (7.27 ± 4.85), and NRAS^WT^ (3.15 ± 3.30) or GFP controls (-1.33 ± 2.85) showed mild antagonism (Fig. S11, S14).

**Figure 6.**
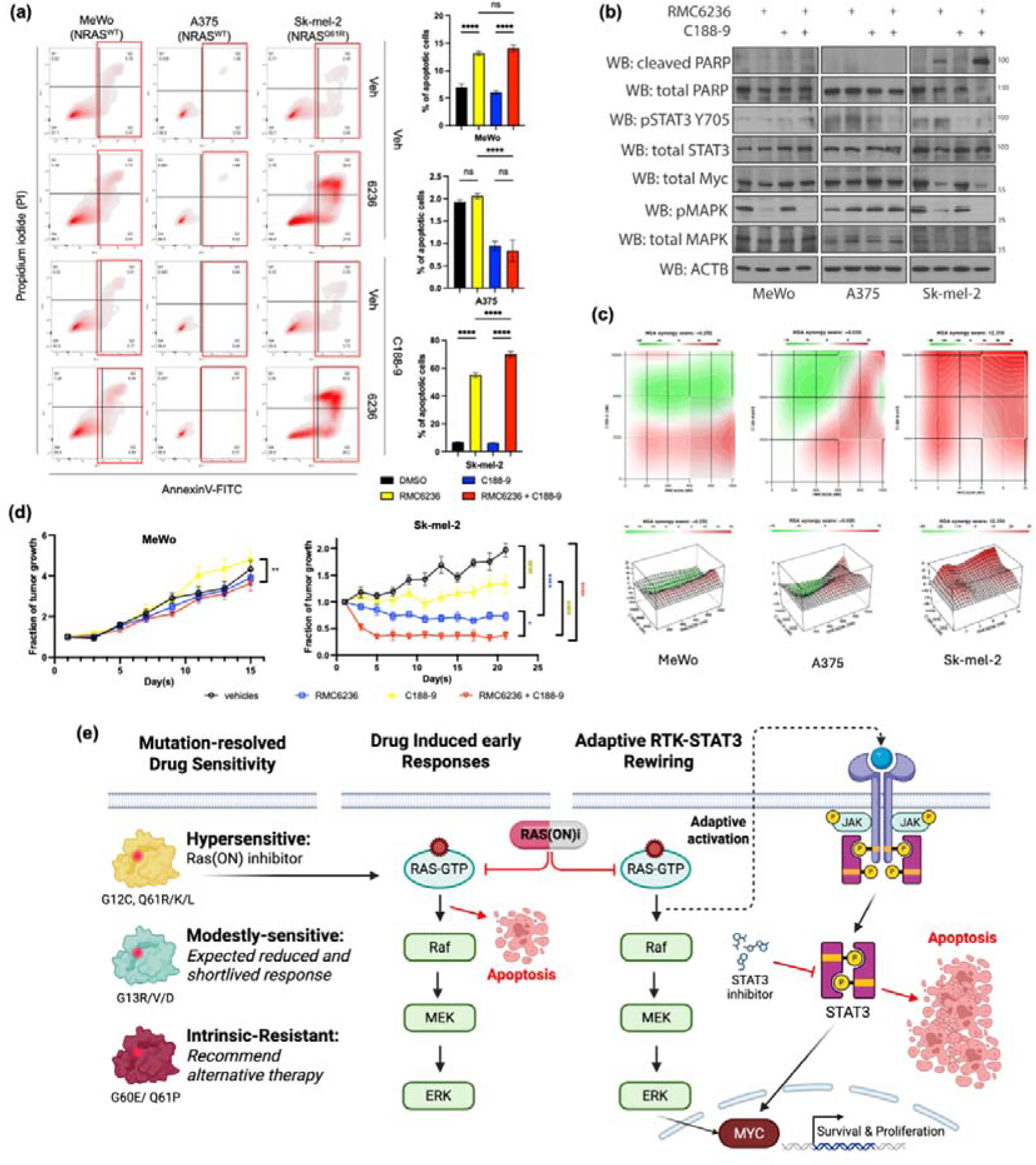
Combined pan-RAS(ON) and STAT3 inhibition exhibits potent antitumor activity in NRAS-mutant melanoma. **(a)** (i) Representative Annexin V-FITC/PI flow cytometry density plots and (ii) quantification of apoptotic cells in NRAS^WT^ (MeWo, A375) and NRAS^Q61R^ (sk-mel-2) cells following 24-hour treatment with RMC-6236 (1 µM for WT; 0.1 µM for Q61R), C188-9 (10 µM), or the combination. Data are displayed as mean ± SD (n=3). Statistical significance was determined by two-way ANOVA followed by a post-hoc multiple comparisons test (***p < 0.001, ****p < 0.0001). **(b)** Western blot analysis of cleaved PARP, p-STAT3 (Tyr705), and MYC levels in melanoma cells after 16-hour treatment, confirming enhanced apoptosis and target suppression upon combination treatment. **(c)** Viability-based synergy matrix of melanoma cells treated with the pan-RAS(ON) inhibitor RMC-6236 and/or the STAT3 inhibitor C188-9. Synergy was quantified using the Highest Single Agent (HSA) model. The combination demonstrates strong synergy in NRAS^Q61R^ (sk-mel-2) cells (HSA score ∼10) compared to moderate synergy in NRAS^WT^ cells (HSA score ∼0). Data are representative of three independent replicates (n=3); drug concentrations are expressed in nM. **(d)** Fractional tumor growth (normalized to Day 1 of treatment) in mice bearing subcutaneous MeWo (NRAS^WT^) or sk-mel-2 (NRAS^Q61R^) xenografts following treatment with vehicle, RMC-6236 (10 mg/kg daily via oral gavage), C188-9 (50 mg/kg daily via oral gavage), or the combination for 15 (MeWo) or 21 (sk-mel-2) days. Data points and bars represent mean ± SEM (n=8 tumors per group). Statistical significance was analyzed by two-way ANOVA followed by post-hoc multiple comparisons tests (*p < 0.05, **p < 0.01, ***p < 0.001, ****p < 0.0001).

Annexin V/PI staining further demonstrated that combining RAS(ON) inhibitors with either C188-9 or napabucasin failed to increase apoptosis in NRAS^WT^ cells (MeWo, A375) beyond monotherapy levels. In contrast, in both endogenous (sk-mel-2) and the OE NRAS mutated models (MeWo NRAS^Q61R/K/L^) showed a significant 1.2–1.5-fold increase in cell death compared to RMC-6236 alone, reaching 70–80% total apoptosis (p < 0.0001) and exhibiting a larger late-apoptotic (Annexin V^+^/PI^+^) population (Fig. 6A; Fig. S10 and S14). Mechanistically, immunoblotting confirmed that while RAS(ON) inhibition alone only partially reduced MYC, its combination with STAT3 inhibitors (C188-9 or napabucasin) triggered near-complete MYC ablation specifically in NRAS-mutant models (MeWo-NRAS^Q61R/K/L^ and sk-mel-2). This profound MYC loss, was not observed in NRAS^WT^ cells, correlated with markedly increased cleaved PARP (Fig. 6B, Fig. S14).

To evaluate therapeutic relevance *in vivo*, we established subcutaneous xenografts and administered vehicle, RMC-6236 (10 mg/kg), or C188-9 (50 mg/kg) as monotherapies or in combination. In MeWo-derived xenografts (NRAS^WT^), tumor growth was comparable across all groups, with volumes increasing approximately 3-fold by the study’s end. Conversely, in sk-mel-2 (NRAS^Q61R^) xenografts, vehicle-treated tumors expanded 1.9-fold, while C188-9 monotherapy achieved stable disease. Notably, RMC-6236 monotherapy induced a 32% tumor shrinkage, whereas the combination treatment resulted in superior tumor shrinkage of 65% compared to the initial volume (Fig. 6D). These results were recapitulated in the MeWo overexpression system using the STAT3 inhibitor napabucasin (10 mg/kg). While NRAS^WT^ tumors showed marginal benefit from the combination (27.3% growth reduction; p < 0.01), NRAS^Q61R^ xenografts demonstrated a robust response, characterized by a ∼57% volume reduction (p < 0.0001) and a significant decrease in tumor weight (53.1%; p < 0.0001). Interestingly, NRAS^Q61R^ xenografts also exhibited a marked loss of pigmentation, appearing pale in contrast to the characteristically melanotic (black) NRAS^WT^ tumors (Fig. S14). Collectively, these *in vivo* data confirm that common Q61 NRAS mutations confer a specific, targetable vulnerability to dual RAS/STAT3 inhibition in melanoma.

## Discussion

RAS mutations are among the most common oncogenic drivers, present in approximately 20–30% of human cancers. KRAS mutations account for about 85% of RAS □ mutated tumors, followed by NRAS (15%) and HRAS (∼1%), with proportions varying by cancer type (26, 27). Although multiple KRAS □ specific inhibitors have now entered clinical use, there has been little progress in the development of NRAS mutation-specific inhibitors. Several therapeutic strategies have been explored for NRAS □ mutant cancers, including NRAS □ targeting antisense oligonucleotides, monobodies designed to destabilize NRAS, and SHOC2 (scaffold protein critical for Q61 □ mutant) inhibitors (13, 28, 29, 30). However, these approaches generally suffer from limitations such as the need for micromolar concentrations to achieve meaningful pathway inhibition, suboptimal potency or isoform/mutant selectivity, and challenges in delivery and intracellular bioavailability, including poor cellular uptake and difficulty crossing biological membranes.

In parallel, substantial advances in broad-spectrum RAS-targeting agents have opened an alternative therapeutic avenue for NRAS-mutated melanoma. Pan-(K)RAS inhibitors such as ADT-007 (14) and BI-2865 (13), as well as tri-complex RAS(ON) inhibitors including RMC-6236 (16) and RMC-7977(17), have demonstrated broad preclinical activity across RAS isoforms and variants. Notably, RMC-6236 (daraxonrasib) has shown encouraging antitumor activity in KRAS-mutated pancreatic and lung cancers in early-phase trials (31) and multiple phase III trials are ongoing (RASolute 301 to RASolute304). Moreover, a recent phase Ib/II study reported objective responses, including one complete and one partial response, in patients with NRAS-mutated melanoma treated with daraxonrasib(19). These advances motivated our systematic evaluation of NRAS variant sensitivities in melanoma.

Overall, our study established the first systematic, mutation-resolved drug-sensitivity map for 95 NRAS missense variants using 3D melanoma spheroids. Distinct from prior NRAS studies, our work integrates saturation mutagenesis, structural modelling, pathway analysis, and *in vitro*/*in vivo* validation to define the functional landscape and drug sensitivities of NRAS-mutated melanoma towards different RAS inhibitors.

As expected, among the six RAS □ targeting agents tested, RMC □ 6236 and RMC □ 7977 demonstrated the broadest and most potent activity, potently suppressing both P □ loop (G12/G13) and Switch II (Q61) mutants. These results strongly support the prioritization of RAS(ON) inhibitors in clinical development, a direction reinforced by translational evidence of their activity across multiple RAS-addicted cancers, including melanoma, lung, colorectal, and pancreatic carcinomas (16, 17, 18, 19, 21). In contrast, the distinct and partially complementary profiles of ADT □ 007 and BI □ 2865 suggest their potential utility against RAS(ON) inhibitor-resistant or less susceptible mutants.

Another major finding is that a subset of oncogenic, recurrent NRAS mutants shows reduced sensitivity to RMC-6236 and RMC-7977, notably NRAS^Q61P^, NRAS^G13D^, NRAS^G13V^, NRAS^G13R^ and NRAS^G60E^. Structural modelling suggests that different mechanisms contribute to this reduced susceptibility. The Q61P and G60E substitutions appear to disrupt essential polar and hydrophobic interactions that support stable formation of the inhibitor-bound tricomplex, as shown in Fig. 3A and Fig. S7. In contrast, the G13D, G13R and G13V substitutions, although located away from the inhibitor binding surface, may weaken the overall stability of the RAS-RAS(ON)-inhibitor-PPIA complex. Consistent with this interpretation, previous studies reported that KRAS G13 mutant cells display higher EC_50_ values toward tricomplex inhibitors compared with other molecular subgroups (17), suggesting that this structural constraint may be shared across multiple RAS isoforms. Although these NRAS variants represent a smaller fraction of NRAS mutated melanoma cases (Table S5), understanding their drug response profiles remains important for guiding treatment selection and patient expectations.

Translational cancer therapy often requires sustained, multi-targeted regimens rather than short-term monotherapy, particularly in aggressive cancers such as melanoma, which frequently develops rapid resistance (23, 32). Recent studies have shown that RAS(ON) inhibitors can remodel the tumor microenvironment in NRAS-mutated melanoma, increasing CD8 □ T-cell infiltration and enhancing responsiveness to immune checkpoint blockade. While this supports the use of immunotherapeutic combinations, not all patients are eligible for such strategies.

Our work therefore explored alternative combination approaches by defining the adaptive state induced by RAS(ON) inhibition. Beyond acute suppression of MAPK output, RMC-6236 and RMC-7977 drove a distinct drug-adapted program characterized by increased STAT3 activity, induction of cytokine-responsive and stress-remodeling transcriptional modules, and enrichment of ligand-, co-receptor-, and RTK-associated signaling features. In parallel, phospho-RTK profiling demonstrated compensatory activation of multiple receptors, including ERBB3, ERBB4, EphA6, MET, DDR1, and RYK. Further pharmacologic interrogation showed that STAT3 reactivation was effectively suppressed by broader kinase inhibitors. Together, these findings support a model in which RAS(ON) inhibition creates a cytokine- and growth factor-rich adaptive state. The inflammatory signaling and redundant RTK inputs converge on STAT3 to preserve melanoma-cell survival under drug pressure. Functionally, genetic or pharmacologic STAT3 inhibition markedly increased apoptosis, enhanced PARP cleavage, drove near-complete MYC loss, and improved antitumor efficacy specifically in NRAS-mutant melanoma models. These data therefore position STAT3 as both a mechanistic mediator of adaptive survival and a rational combination target for deepening and prolonging responses to tri-complex RAS(ON) inhibitors (Figure 6).

Evidence from other RAS-driven cancers reinforces the importance of co-targeting adaptive signaling nodes when oncogenic RAS is inhibited. In KRAS G12C non-small cell lung cancer, KRAS inhibition rapidly induces STAT3 activation, and combined inhibition of KRAS and STAT3 improves tumor control and enhances natural killer cell cytotoxicity (32). Additional studies show that co-inhibition of FLT3 (20), CDK4/6 (33), SHP2/MEK (17), or MAPK/PI3K(22) can augment the activity of RMC-6236 and RMC-7977 across multiple RAS mutant models indicating that effective suppression of compensatory signaling is essential for durable responses.

Our findings converge to offer actionable insights for clinical translation and clinical trial design. The broad sensitivity profile we observed suggests that RAS(ON) inhibitors should serve as a backbone therapy for the majority (∼95%) of NRAS-mutated melanomas. For the subset of patients harboring moderately sensitive or resistant variants (e.g., Q61P, G13D/V/R, G60E), mutation-specific exclusion or alternative strategies (such as ADT-007 or BI-2865) may be necessary. Furthermore, the adaptive STAT3 activation identified here provides a mechanistic rationale for combination regimens to enhance the depth and durability of RAS inhibition responses. This approach to rational combination therapy aligns with the broader paradigm of integrating RAS inhibition with other targeted or immunomodulatory agents across diverse cancers (11, 13, 34, 35). While our phospho-RTK array and inhibitor studies support a convergent multi-RTK input into STAT3 reactivation, do not yet assign a single dominant upstream receptor. Future studies incorporating endogenous perturbation of candidate RTKs, patient-derived models, and on-treatment clinical specimens will be important to refine the receptor hierarchy and identify biomarkers of response to combined RAS(ON)/STAT3 inhibition.

Furthermore, our saturation mutagenesis screen identified NRAS variants that, while not currently classified as clinically recurrent, hold potential for future emergence. The resulting drug sensitivity atlas therefore serves as a proactive, predictive resource to inform clinical decision-making should these mutations appear in patient genomic profiles.

In summary, this mutational-drug sensitivity atlas defines the intrinsic vulnerabilities and adaptive survival mechanisms of NRAS-mutated melanoma under direct RAS inhibition. Our findings define mutation-encoded and adaptive response layers in NRAS-mutated melanoma, supporting mutation-guided precision use of RAS(ON) inhibitors and rational STAT3-based combination regimens to improve efficacy.

## Ethics Compliance Statement

The animal study protocol was approved by the Animal Experimentation Ethics Committee of the Chinese University of Hong Kong (reference number: 22-257-MIS). All animal experiments were performed in accordance with the Animals (Control of Experiments) Ordinance of Hong Kong SAR, China, and complied with all institutional and national guidelines for the care and use of laboratory animals. Analyses of human-derived genomic and clinical data (TGEN, Genome Res 2017; MSK, Clin Cancer Res 2021; DFCI, Nat Med 2019; COSMIC) were conducted using publicly available datasets. In relation to tumor size, we confirm the following: (i) The maximal tumor size permitted by the CUHK Animal Experimentation Ethics Committee is 1.5 cm in the longest diameter or 1.5 cm³ in volume. (ii) This limit was not exceeded in any experimental animal. Throughout the study, tumor dimensions were measured every day (or every 2 days, depending on experimental aims),and mice were euthanised immediately if tumors approached the approved threshold or if signs of distress were observed. No animal developed ulceration, impaired mobility, or tumor burden exceeding ethical guidelines.

## Data Availability Statement

Data from combined melanoma cohorts, including TGEN, Genome Res 2017, MSK, Clin Cancer Res 2021, and DFCI, Nat Med 2019, are publicly available via cBioPortal (www.cbioportal.org; accessed June 20, 2024). COSMIC data were retrieved from the COSMIC database for August 2025. The data generated in this study are provided in the article and Supplementary Materials. Original datasets, additional materials, codes, parameter files, and raw data are available from the corresponding author upon reasonable request.

## Acknowledgments

We thank Professor Hui Zhao for kindly providing Ba/F3 cells. We also acknowledge the core facility of the School of Biomedical Sciences, Chinese University of Hong Kong, for technical assistance and support. We also thank Professor Carol Tong Man for her insightful comments and supportive guidance during the refinement of this manuscript. The figures were created at https://BioRender.com.

## Supplementary Materials

Detailed methods, additional figures, and data will be provided online as a part of a published article. These materials are accessible to all readers and provide essential supporting information that complements the main text.

## Author Contributions

Conceptualization: S.F.Y., C.C.; Methodology: T.Y.L., W.Y., X.C., S.F.Y., C.H.L., M.F.L., K.L., M.T., B.K.B, P.K.C., C.N.L.; Validation: X.C., S.F.Y., W.Y.; Formal analysis: W.Y., X.C., S.F.Y., T.Y.L.; Data Curation and Investigation: C.L., C.H.L., X.C., S.F.Y.; Resources: S.K.W.T., C.C., K.K.L.C.; Writing—original draft preparation: S.F.Y., X.C. ;Writing—review and editing: S.F.Y., T.Y.L., X.C., K.L., C.C., K.Y.S.,; Visualization: S.F.Y., X.C., T.Y.L.; Supervision: S.K.W.T.; Project administration: S.F.Y., K.L., S.K.W.T.; Funding acquisition: S.K.W.T., K.K.L.C., B.K.B..All authors reviewed and approved the final manuscript.

